# High-throughput Single-cell Proteomics Enabled by an Integrated Hyperplexing and Automatic Labelling Approach without Fractionation

**DOI:** 10.1101/2025.06.22.660561

**Authors:** Hui Zhang, Qing Zeng, Kai Jin, Chuanxi Huang, Yi Liu, Xing Liu, Jing Yang, Hanbin Ma, Fuchu He, Yun Yang

## Abstract

Single-cell proteomics (scProteomics) enables comprehensive analysis of protein composition, expression and functions at the single-cell level. While label-free techniques have generated promising results, achieving high-throughput scProteomics remains a substantial challenge. In this study, we present an approach that increases scProteomics throughput by approximately 30-fold through the true integration of isobaric tags (IBT) with tandem mass tags (TMT), a method we term ‘Integral-Hyperplex’. Due to remarkable differences in signal response between TMT16 and IBT16 reporter ions, we implemented two normalization strategies to ensure accurate and integrated quantification: (1) the inclusion of a normalization channel in both IBT and TMT groups to serve as internal standards; and (2) the generation of protein-specific conversion factors between IBT16 and TMT16 signals. We validated the quantification accuracy of the Integral-Hyperplex method by labeling HeLa digests at varying ratios. Furthermore, we demonstrated fully automated labeling on the active-matrix digital microfluidics chip, consuming only 12 nL of labeling reagent per single cell and with a reaction volume as low as 20 nL. Our approach was validated through proof-of-principle quantification experiments across three types of single cells. Approximately 2,000 protein groups were quantified per cell, and the three cell types can be well clustered, confirming the reliability of the Integral-Hyperplex approach. Approximately 300 samples can be analyzed per day due to insufficient resolution of timsTOF SCP. Throughput can be further enhanced using TMT32, additional labeling reagents, and more advanced mass spectrometers such as Orbitrap Astral and timsTOF Ultra 2, and our approach is theoretically capable of analyzing up to ∼2,000 single cells per day.

## INTRODUCTION

Single-cell analysis provides critical insights into cell populations, differentiation, function, and unique features of cell subtypes within complex cell systems.^1,2^ In-depth analysis of single cells reveals subtle differences between cells that are often overlooked in traditional population-level analysis.^3^ Single-cell proteomics (scProteomics) aims to comprehensively characterize protein expression at the single-cell level without bias, thereby uncovering biological functions and intercellular heterogeneity.^4^ Despite recent progress, scProteomics still faces technical challenges, including limited proteome coverage, low throughput, and insufficient robustness. Mass spectrometry (MS)-based scProteomics has advanced rapidly, with many label-free approaches developed in recent years, such as nanoliter scale oil-air-droplet (OAD) chip^5^, nanodroplet processing in one pot for trace samples (nanoPOTS)^6^, integrated device for single-cell analysis (iPAD-1)^7^, true single-cell-derived proteomics (T-SCP)^8^. While these label-free methods have yielded promising results, their low throughput in MS detection remains a major bottleneck—especially for large-scale applications.

To address this limitation, multiplexing strategies such as isobaric labeling have been employed to increase throughput in scProteomics. As first demonstrated in the Single Cell ProtEomics by Mass Spectrometry (SCoPE-MS) workflow,^9^ isobaric labeling has become a widely adopted technique. Subsequent developments in recent years include nanoPOTS combined with TMT 10-plex (TMT10)^10^, nested nanoPOTS (N2) chip combined with TMT10^11^, SCoPE2^12^ and Single Cell pRotEomE aNalysis (SCREEN)^13^. To further improve the throughput in MS detection, a few hyperplexing approaches have been reported in scProteomics. Most current methods rely on a single TMT kit, where increasing the number of channels requires ultra-high MS resolution to distinguish subtle mass differences. Alternatively, combining multiple labelling reagents in a single run (hyperplexing) offered another solution. Liang et al. developed the HyperSCP method, which combined 2-plex SILAC (stable isotope labelling of amino acids in cell culture) isotopic labelling with TMTpro 18-plex (TMT18) isobaric labelling, enabling the analysis of up to 28 single cells in a single LC-MS run.^14^ To date, however, hyperplexing approaches that combine distinct isobaric kits have only been applied to bulk samples. Multiplexing levels of 27-plex^15^ and 29-plex^16^ have been achieved by combining TMT 11-plex (TMT11) with TMT 16-plex (TMT16) or TMT18 reagents. Wu et al. further achieved throughput to 45-plex by combining IBT16, TMT11, and TMT18 in a single run.^17^ However, the shared 11 channels between TMT11 and TMT18 compromised quantitative accuracy in those channels. Wu et al. later extended this method to 102-plex^18^ by introducing alanine or glycine residues to peptides prior to IBT16 or TMT18 labelling. Nevertheless, these samples were typically fractionated into 20 to 40 segments, limiting the overall throughput improvement to only 2–3-fold. Meanwhile, multiplexed single-cell proteomics sample preparation still faces challenges in handling minute sample volumes, multi-step reactions, and reagent consumption. Digital microfluidics (DMF), a novel droplet-based manipulation technology^19^, has therefore emerged as a promising solution for automated and nanoliter-scale single-cell sample processing^20,21^.

In this study, we developed a high-throughput method by integrating IBT 16-plex and TMT16 in a single run without fractionation. To overcome the remarkable signal discrepancies between IBT16 and TMT16, we introduced two normalization strategies to enable accurate and integrated quantification. It is noteworthy that despite improvements in mass spectrometry throughput, the sample preparation process still faces challenges such as complex operations and protein adsorption loss, which limit overall analytical efficiency and reproducibility. To address this, we integrated the BOXmini™ SCP instrument to achieve fully automated sample labeling on the AM-DMF chip.^22^ Using this high-throughput strategy, we achieved deep and accurate proteome coverage in individual 293T, HeLa, and A549 cells.

## EXPERIMENTAL SECTION

### Quantitative accuracy evaluation of TMT and IBT labelling

HeLa digest with varying protein amounts were prepared in a fixed volume. For ratios of 1:2:3:4:1:2:3:4, 0.5, 1.0, 1.5, 2.0, 0.5, 1.0, 1.5 or 2.0 ng/μL HeLa digest were labelled with either TMT16 or IBT16 channel (5 μg/μL) in 25 mM HEPES for 2 h. For ratios of 1:2:3:4:5:6:7:8, 0.5, 1.0, 1.5, 2.0, 2.5, 3.0, 2.5 or 4.0 ng/μL HeLa digest were labelled with TMT16 or IBT16 in the same way. After quenching with 5% hydroxylamine, the 16 labelled aliquots (8 TMT and 8 IBT channels) were combined and dried using a SpeedVac. Labelled peptides were redissolved in 0.1% FA and 5 ng of HeLa digest in total was injected each time.

To evaluate quantitative accuracy at the single-cell-equivalent level, experimental ratios were set to 1:2:3:1:2:3. 100, 200, 300, 100, 200, and 300 pg/μL HeLa digest was labelled with TMT16 or IBT16 (5 μg/μL), followed by quenching with 5% hydroxylamine. The samples labelled with 6 TMT or 6 IBT channels were then combined, and carrier peptides at 25x (5 ng), 50x (10 ng), or 100x (20 ng) levels were directly added prior to nanoLC-MS analysis.

### Normalization between IBT and TMT reporter ions

The single-cell equivalent dataset at the level, with experimental ratios set to 1:2:3:1:2:3 and 50x carrier, was used for normalization test. The first method is to add one normalization channel in both IBT and TMT groups, and the first 1x channels in IBT and TMT groups were designated as the normalization channel in this study. The same sample was labelled by IBT or TMT, and protein intensities were normalized by dividing by the corresponding normalization channel value. The second method constructed protein-specific conversion factors by calculating the IBT/TMT ratio for each protein across corresponding IBT and TMT channels (e.g., 1x IBT and 1x TMT). Ratios 1–6 were calculated for the six TMT channels (1:2:3:1:2:3), and the mean IBT/TMT ratio was used as a conversion factor for each protein. Final intensities were adjusted using: ‘TMT value × conversion factor’.

### Manual labelling of single cells

In manual labelling, single 293T cells were isolated and pretreated using the cellenONE system, as previously described under ‘Optimization of TMT and IBT Labelling Conditions.’ TMT16 and IBT16 reagents, pre-dissolved in anhydrous acetonitrile (5 μg/μL), was added to each well at room temperature. The reaction proceeded for 2 h at 1,000 rpm. Unreacted reagents were quenched with 0.6 μL of 5% hydroxylamine. Labelled cell digests were manually combined, followed by addition of carrier and normalization peptides. Finally, samples were transferred to autosampler vials (Waters, QuanRecovery) for nanoLC-MS analysis.

### Automated labelling of single cells using BOXmini™ SCP

The entire automated labelling process (Movie S1, Supporting Information) was conducted within an active matrix digital microfluidic (AM-DMF) chip (model AM16K)^22^, which was operated using the BOXmini™ SCP instrument. The BOXmini™ SCP instrument, the chip, and the associated software were provided by ACX Instruments (Cambridge, UK) and ACXEL Micro & Nano Tech (Foshan, China). The chip consists of 16,384 electrodes arranged in a 128 × 128 array, with an electrode pitch of 250 μm and a gap size of 50 μm between the two plates. The internal environment of the AM16K chip was filled with the medium oil hexadecane, and each sample was wrapped by this oil phase environment, which effectively reduced evaporation and minimized cross-contamination.^22^ The lysis buffer consisted of 0.1% n-Dodecyl-β-D-Maltoside (DDM), 10 mM Tris(2-carboxyethyl) phosphine (TCEP), 40 mM Chloracetamide (CAA), 100 mM 2-[4-(2-hydroxyethyl) piperazin-1-yl]-ethanesulfonic acid (HEPES) and 0.1% RapiGest. Specific cell pretreatment steps prior to labelling followed previously reported protocols.^22^ Briefly, cells were washed over three times with ice-cold PBS and diluted in PBS plus 0.05% Pluronic F68 to a concentration of ∼ 4.0 × 10^5^ cells/mL. Droplets containing a single cell were selected for reduction and alkylation at 56 °C for 15 min, followed by digestion at 37 °C for 2 h. Next, 0.2 μL of 4 mg/mL labelling reagent (in 80% acetonitrile and 100 mM HEPES) was loaded into the chip, and the labelling reaction was carried out at 25 °C for 1 h. The labelling reaction was terminated by adding a quenching reagent (0.5% HA solution in 100 mM HEPES) at 25 °C for 15 min. Four or two labelled cell digests with different labels were combined and diluted with 0.1% formic acid (FA). The samples were subsequently transferred into autosampler vials (Waters, QuanRecovery) using 100 µm I.D. capillary tubes. Labelled samples from three AM16K chips were pooled to obtain a 10-plex sample. Carrier (10 ng/μL, IBT-121C and TMT-133C, or IBT-122 and TMT-134N) and normalization channel (0.5 ng/μL, IBT-114 and TMT-126, or IBT-115N and TMT-127N) samples were then added manually.

### Data Analysis

The raw data were analyzed using PEAKS Online software against the human UniProt FASTA database (20,375 entries). The precursor mass error tolerance was set to 15 ppm and the fragment mass error tolerance was set to 0.05 Da. Enzyme specificity was set to Trypsin/P with up to two missed cleavages allowed. The TMT/iTRAQ Label Quantification mode in the PEAKS Q (de novo-assisted quantification) module was selected for protein quantification. Oxidation (+15.995 Da) on methionine residues and protein N-terminal acetylation (+42.011 Da) were set as variable modifications, while TMT16 (+304.207 Da) or IBT16 (+277.188 Da) were used as fixed modifications. The PEAKS DB (in-depth de novo-assisted search) module was used to calculate labelling efficiencies, where oxidation (+15.995 Da) on methionine residues, protein N-terminal acetylation (+42.011 Da), and TMT16 (+304.207 Da) or IBT16 (+277.188 Da) were selected as variable modifications.

## RESULTS AND DISCUSSION

### Design of the TMT-IBT strategy

In scProteomics, the ever-increasing number of single cells to be analyzed has heightened the demand for high-throughput MS detection. In our strategy, we integrated commercially available IBT16 and TMT16 labelling reagents into a single run. Since IBT16 and TMT16 generate distinct reporter ions and have no shared channels, our approach reduces interference and enhances quantitative accuracy. Importantly, to improve MS detection throughput, no fractionation was applied.

In this study, due to the resolution limitations of the timsTOF SCP, reporter ions from neighboring channels cannot be well differentiated. Therefore, we divided all the 32-plex channels into two groups, ensuring that neighboring channels from TMT16 and IBT16 were assigned to separate groups (Figure 1a). As a result, Group A included 114, 115C, 116C, 117C, 118C, 119C, 120C, and 121C from IBT16, and 126, 127C, 128C, 129C, 130C, 131C, 132C, and 133C from TMT16; while Group B consisted of 115N, 116N, 117N, 118N, 119N, 120N, 121N, and 122 from IBT16, along with 127N, 128N, 129N, 130N, 131N, 132N, 133N, and 134N from TMT16.

**Figure 1.**
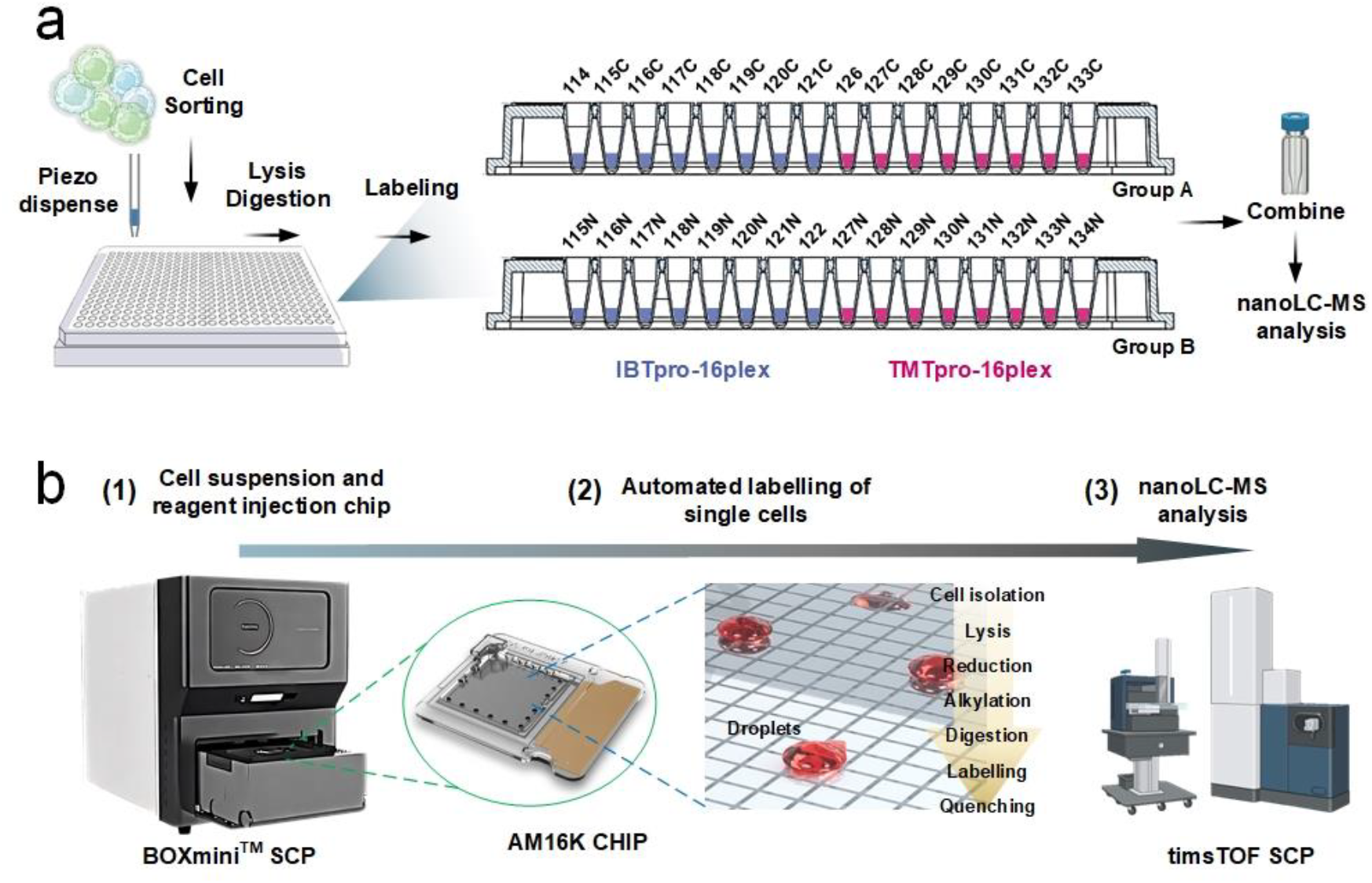
Schematic diagram of our TMT-IBT hyperplexing strategy. (a) Workflow for the manual TMT-IBT hyperplexing strategy. (b) Workflow for the automated TMT-IBT hyperplexing strategy using the BOXmini™ SCP platform. This workflow involves loading cell suspension, lysis, digestion, and labelling reagents onto the chip, followed by execution of a pre-programmed sample processing protocol. Finally, the pooled samples are transferred to an autosampler vial for nanoLC-MS/MS analysis. (c) The expanded view of the layout and major reactions on the AM16K chip.

For the manual labelling process, we used the cellenONE instrument to precisely isolate single cells and dispense them into 384-well plates. After a one-step pretreatment to obtain peptides, labelling and quenching reagents were sequentially added to the wells. The labelled samples were manually pooled into Group A or B for nanoLC-MS/MS analysis. Throughout the process, only the pooled samples required transfer from the wells into autosampler vials.

To automate the TMT-IBT labelling of single cells, we used the recently released BOXmini^TM^ SCP instrument, in which all reactions were performed on an AM16K chip (Figures 1b and S1). Parallel sample processing for multiple cells was achieved, and the entire sample preparation workflow—including single-cell isolation, lysis, protein reduction, alkylation, enzymatic digestion, labelling, and quenching—was fully integrated within this digital microfluidic chip. Moreover, each droplet volume was merely ∼3.125 nL, with total reaction volumes under 20 nL, facilitating efficient reaction kinetics and minimizing reagent consumption. This ultra-low volume also limited the use of labelling reagents to just 20 nL per reaction, minimizing labelling costs.

### Optimization of TMT Labelling and IBT Labelling conditions

Utilizing commercially available HeLa digest, labelling conditions for TMT0 and IBT-2 (115N, 118C) were optimized, favored for their cost-effectiveness. The labelling procedure was initially evaluated using 25 ng of HeLa digest. As shown in Figure 2a, labelling efficiency increased with the amount of reagent used, although the total number of proteins identified improved only marginally. From a cost-benefit perspective, lower amounts of TMT reagent which can achieve satisfactory labelling efficiency is optimal. To optimize IBT labelling (Figure 2b), we similarly examined how reagent dosage affected protein identification and labelling efficiency. IBT-115N and IBT-118C were tested in varying amounts. Considering reagent cost, labelling efficiency, and proteome coverage, 5.0 µg was chosen as the optimal dosage to obtain sufficient results. We further optimized the procedure by evaluating different buffer systems and concentrations. As shown in Figure 2c, using 40 mM TEAB as a buffer resulted in lower protein identification and labelling efficiency compared to 40 mM HEPES. Interestingly, labelling efficiency improved and more proteins were quantified when the HEPES concentration was reduced to 25 mM, whereas higher HEPES levels appeared to inhibit the reaction. Next, we optimized the amounts of TMT0 and IBT-115N labelling reagents for single HeLa cells using 25 mM HEPES buffer. Both reagents achieved over 95% labelling efficiency and highest number of quantified proteins at dosage of 7.5 µg, (Figure 2d). Based on these results, a standardized protocol for manual labelling of cell digests and single cells was established using the optimized conditions.

**Figure 2.**
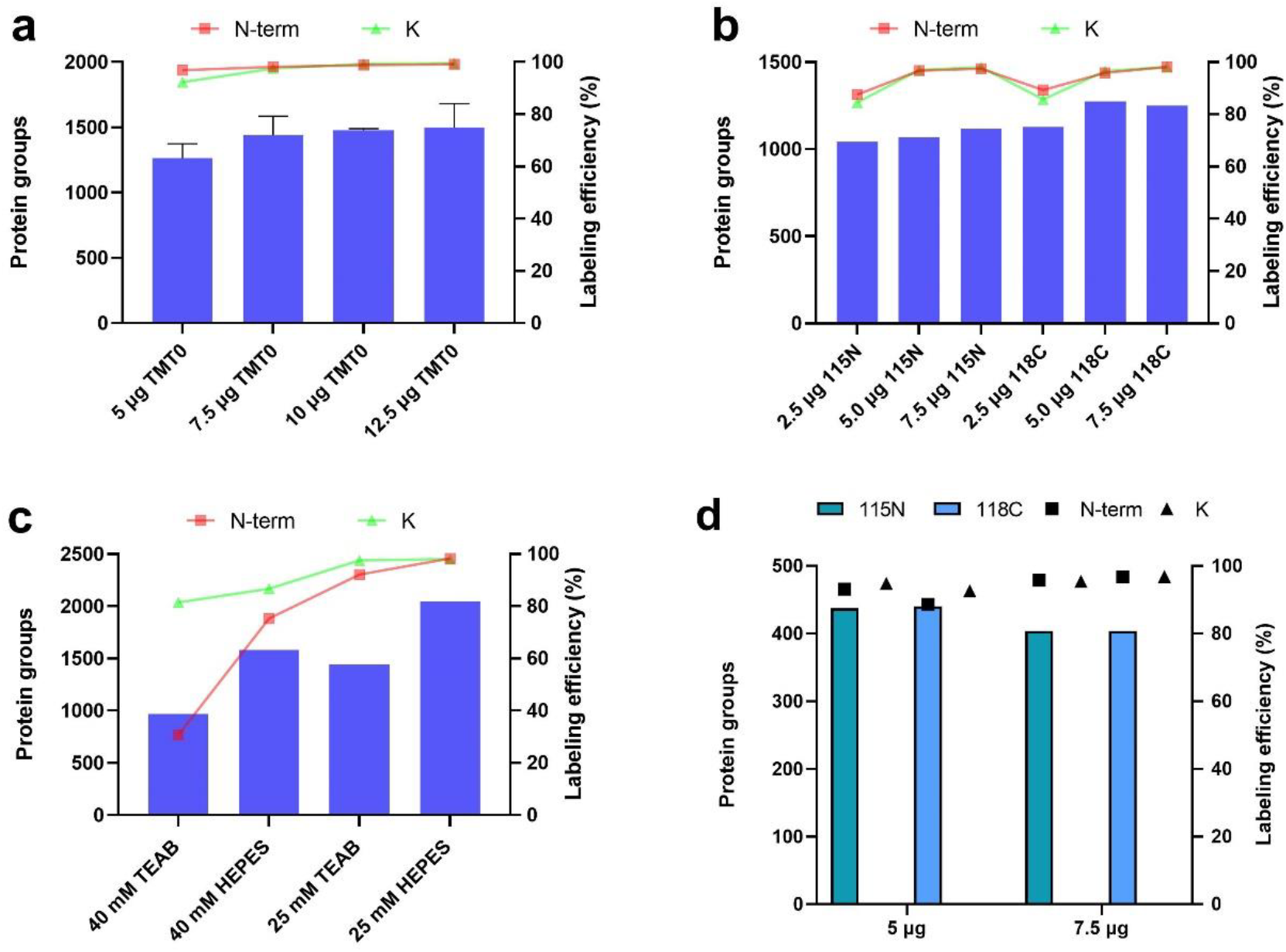
Optimization of TMT and IBT labelling conditions. Optimization was based on both proteome depth and labelling efficiency, with 1 ng of labelled HeLa digest injected per run. (a) Dosage optimization of the TMT0 reagent using 25 ng of HeLa digest. (b) Dosage optimization of IBT-115N and IBT-118C using 25 ng of HeLa digest. (c) Comparison of different concentrations of TEAB and HEPES. (d) Dosage optimization for single HeLa cells.

### Quantitative accuracy evaluation in separate TMT or IBT labelling

Since single cells typically contain only pico-to nanogram levels of protein, the quantitative accuracy of TMT and IBT labelling was initially assessed at the nanogram level. HeLa digest samples with varying protein amounts were combined in a fixed volume, and 5 ng of the digest was injected. The protein amounts in different TMT or IBT channels were mixed in known ratios of 1:2:3:4:1:2:3:4 or 1:2:3:4:5:6:7:8. As shown in Figures 3a-3b and S2a-b, the resulting ratios were close to the true ratios in both TMT and IBT channels. Meanwhile, good repeatability was observed for the 1:2:3:4 ratios in both TMT and IBT channels. Both groups showed some deviation, indicating distinct response patterns between the TMT and IBT channels, posing a great challenge for quantitative integration of TMT and IBT channels. When the ratios were extended to 1:2:3:4:5:6:7:8, as shown in Figures 3c-3d, a similar trend was observed. The resulting ratios demonstrated similar accuracy and precision in both TMT and IBT channels compared to the 1:2:3:4:1:2:3:4 ratios. Overall, TMT channels exhibited better quantification accuracy than IBT channels.

**Figure 3.**
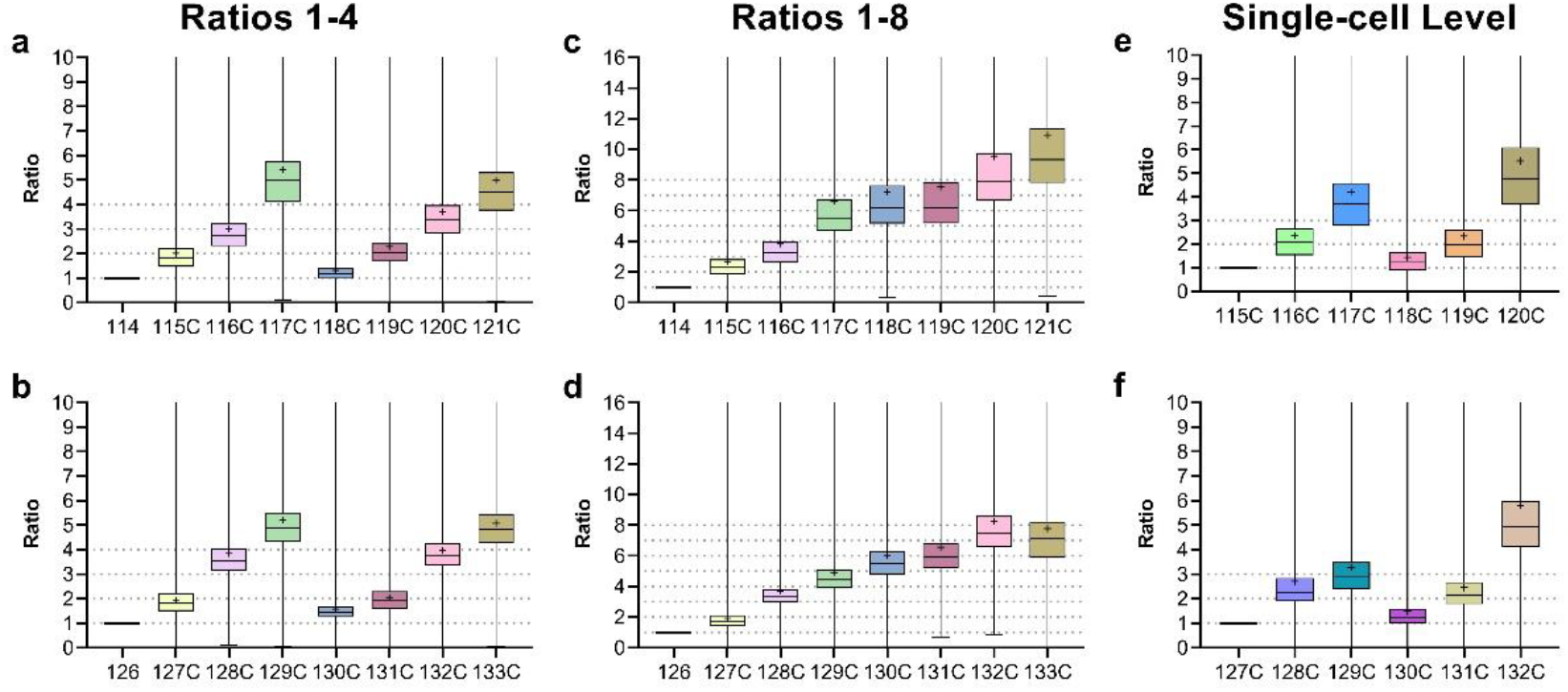
Evaluation of the quantitative accuracy of TMT and IBT labelling. (a–b) Observed ratios in IBT16 (a) and TMT16 (b) Group A, with a preset ratio of 1:2:3:4:1:2:3:4 (1 = 1 ng). (c–d) Observed ratios in IBT16 (c) and TMT16 (d) Group A, with a preset ratio of 1:2:3:4:5:6:7:8 (1 = 1 ng). (e–f) Observed ratios in IBT16 (e) and TMT16 (f) Group A, with a preset ratio of 1:2:3:1:2:3 (1 = 200 pg) and 50× carrier. The first channel in each group was left blank and is not displayed. Protein group ratios are plotted as box plots, with expected ratios indicated by dashed lines. Boxes depict the interquartile range, ‘+’ symbols represent mean values, and central bands indicate medians.

When evaluating the quantitative accuracy at the single-cell equivalent level, experimental ratios were set to 1:2:3:1:2:3, with 1 representing 200 pg. A carrier channel was introduced to boost MS1 signals for each sample channel, thereby increasing the number of identified proteins. As shown in Figures 3e-3f, where a 50x carrier channel was added, high quantification accuracy was achieved at the single-cell equivalent level, except for the 120C and 132C channels adjacent to the carrier channels. These adjacent channels were excluded in subsequent TMT-IBT experiments for single cells. Additionally, the quantitative accuracy was compared for carrier amounts of 100x (Figure S3a), 50x (Figures 3e-f and S2e-f), and 25x (Figure S3b). The number of identified proteins increased with the carrier amount; however, a 100x carrier decreased quantitative accuracy. Therefore, a 50x carrier was selected as the optimal condition. Under these conditions, good quantitative accuracy was achieved at the single-cell level. Overall, good quantitative accuracy and repeatability were observed for the labelled single-cell equivalents.

### Normalization between IBT and TMT reporter ions

In the literature, quantitative results have typically been analyzed separately, and integrated IBT and TMT labelling has not been reported, due to the substantially different responses of IBT and TMT reporter ions in MS detection. Although accurate quantification was achieved in separate TMT or IBT labelling experiments (Figures 3e-f and S2e-f), direct quantitative comparisons between IBT and TMT groups resulted in poor quantification accuracy (Figures 4a-b), and vice versa. Specifically, the quantitative differences for each protein or peptide between IBT and TMT groups were highly irregular and diverse (Figure S4), making it impossible to normalize them using a uniform formula applicable to all proteins. Consequently, each protein requires individual normalization.

**Figure 4.**
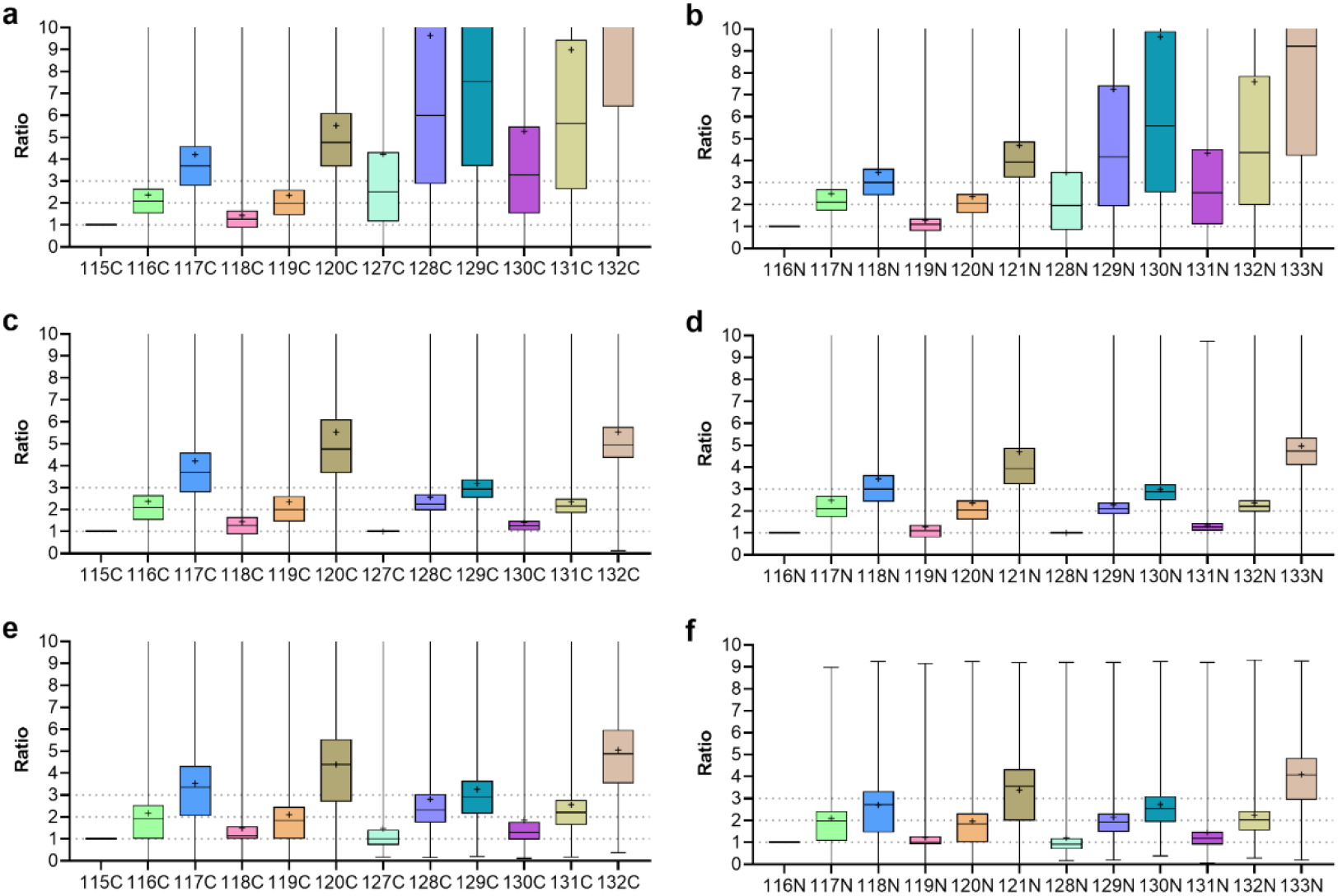
Normalization of reporter ion intensities between IBT16 and TMT16. (a–b) Observed ratios in TMT-IBT Group A (a) and Group B (b) for the preset ratio of 1:2:3:1:2:3, before normalization. (c–d) Box plots showing protein group ratios normalized using normalization channels in Group A (c) and Group B (d). (e–f) Box plots showing ratios normalized using conversion factors for Group A (e) and Group B (f). ‘+’ symbols indicate mean values, and ‘—’ symbols indicate median values.

To address this issue, we developed two normalization strategies specifically designed to correct the intensity differences for each protein labelled by IBT16 or TMT16. The first method introduces a normalization channel in both the IBT and TMT groups, in which the same sample is labelled by IBT or TMT and serves as a ‘normalization standard’. Normalized signals for each protein in other channels are calculated by dividing their protein intensities by that of the normalization channel. The normalized intensities of IBT and TMT can then be directly compared for relative quantification, and satisfactory results were achieved as shown in Figures 4c-d, using 115C and 127C, and 116N and 128N as normalization channels, respectively. Furthermore, these normalization channels can also be used to normalize samples across multiple batches, helping to reduce batch effects.

The second method involves constructing protein-specific conversion factors (Figures 4e-f) by calculating the IBT/TMT ratio for each co-quantified protein across all sample channels. After averaging these ratios across multiple channels, the mean IBT/TMT ratio is used as a conversion factor for each protein in all TMT channels. This factor is then applied to convert the quantitative values of co-quantified proteins in TMT channels using the formula: ‘TMT value × conversion factor’. Quantification accuracy was significantly improved using this approach compared with the unnormalized data (Figures 4a-b). Compared to the ‘normalization channel’ method, the conversion factor approach offers higher throughput but is more complex and generally provides lower normalization accuracy. Both methods achieved effective normalization and successfully mitigated quantification bias.

### Manual TMT-IBT labelling of single cells

To apply our integrated TMT-IBT labelling strategy to single cells, we first tested manual labelling of single 293T cells that had been isolated and pretreated using the cellenONE system. Following TMT-IBT labelling, approximately 1,500 protein groups were quantified from individual 293T cells (Figures 5a-b, Table S2). This proteome depth exceeds that of the recently published high-throughput nPOP (nano-proteomic sample preparation) method^24^, which reported ∼1,000 proteins per human cell using TMT Pro 32-plex. Notably, TMT channels quantified significantly more protein groups than IBT channels, indicating a better response of TMT reporter ions compared to IBT reporter ions on the timsTOF MS platform, which may differ from the behavior observed in bulk samples analyzed on Orbitrap MS.^17,18^

**Figure 5.**
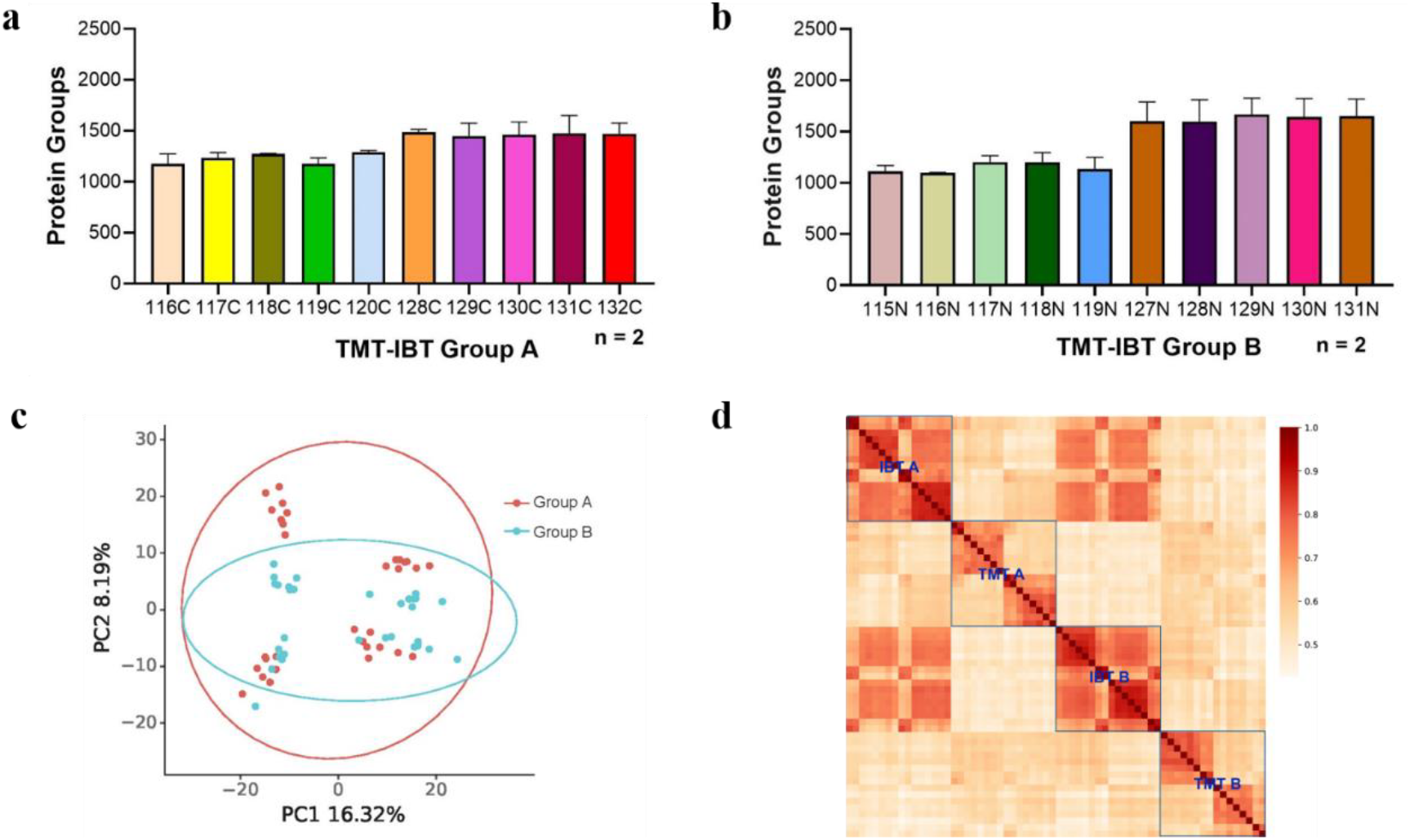
Results of manually labelled single 293T cells. (a–b) Number of identified proteins in TMT-IBT Group A (a) and Group B (b). (c) PCA analysis of Groups A and B. (d) Pearson correlation analysis among four groups: IBT Groups A and B, and TMT Groups A and B.

In addition, cluster analysis was performed, and 293T cells in groups A and B cannot be distinguished, indicating high similarity in these two groups (Figure 5c). Moderate-to-high correlations were observed among the manually labelled single cells (Figure 5d), reflecting cellular heterogeneity. However, distinct differences were observed between IBT and TMT groups and between groups A and B, clearly proving the demand to normalize TMT and IBT signals. Furthermore, labelling efficiencies reached only ∼85-93% in this manual TMT-IBT labelling test, possibly due to the low reagent concentrations used in relatively large reaction volumes and insufficient buffer concentrations to maintain pH ranges during labelling reactions. This limitation needs to be addressed prior to broader applications. While increasing reagent concentrations could improve efficiency, it would also raise costs and introduce excessive salt contamination. As an alternative, reducing reaction volumes and the amount of salts through the use of microfluidic devices may provide a more practical solution.

### Automated TMT-IBT labelling of single cells on BOXmini^TM^ SCP

To further improve labelling efficiency and stability, we utilized the recently developed BOXmini™ SCP instrument in this study. The BOXmini™ SCP has demonstrated advantages in label-free scProteomics;^22^ however, its performance in multiplexed applications has not yet been evaluated. Multiplexed sample preparation for single-cell proteomics remains challenging due to its complexity and labor-intensive nature. Key considerations include ensuring complete transfer of single-cell proteins into the mass spectrometer and enhancing automation to improve sensitivity and throughput.

Here, we assessed an automated workflow for preparing TMT-IBT-labelled single-cell samples using the BOXmini™ SCP platform. The BOXmini™ SCP is equipped with an AM16K chip, which integrates the entire sample preparation process, including single-cell screening, lysis, enzymatic digestion, labelling, and reaction termination— all within a single chip. The chip’s water-in-oil environment prevents evaporation of nanoliter-scale volumes, allowing total reaction volumes as low as 20 nL. This setup minimizes protein loss and ensures high reaction efficiency. Furthermore, only ∼12 nL of labelling reagent is required per cell, significantly reducing reagent consumption and thereby minimizing costs to ∼ 0.47 US Dollar/cell.

By automating our integrated TMT-IBT strategy for 293T, HeLa, and A549 cells on the BOXmini™ SCP, labelling efficiencies increased to over 95% across all cell types (Figure S6), indicating that reduced reaction volumes improve labelling performance. The average number of protein groups quantified per cell ranged from 1,500 to 2,000— approximately 500 more than with manual labelling (Figure 6a). Variability in the number of quantified protein groups also decreased significantly (Figure S7), reflecting the high stability of automated operations on the BOXmini™ SCP. Importantly, dynamic ranges of quantified protein groups from single cells achieved five orders of magnitude (Figure 6b). Using normalized intensities from normalization channels, 100 single cells of each cell type were automatically and accurately separated by principal component analysis (PCA) according to cell type but not TMT or IBT labelling, which indicates our strategy successfully mitigated intensity differences between TMT16 and IBT16 reporter ions (Figure 6c). Importantly, the discrimination was comparable or even better than recently published label-free results of three cell types^22,25^. Correlations within the same cell type were much higher than those between different cell types (Figure 6d), and mean Pearson correlation coefficients were 0.785, 0.798 and 0.725 for 293T, A549 and HeLa cells, respectively;Distinct protein expression patterns across those three human cell lines were evident (Figure 6e).

**Figure 6.**
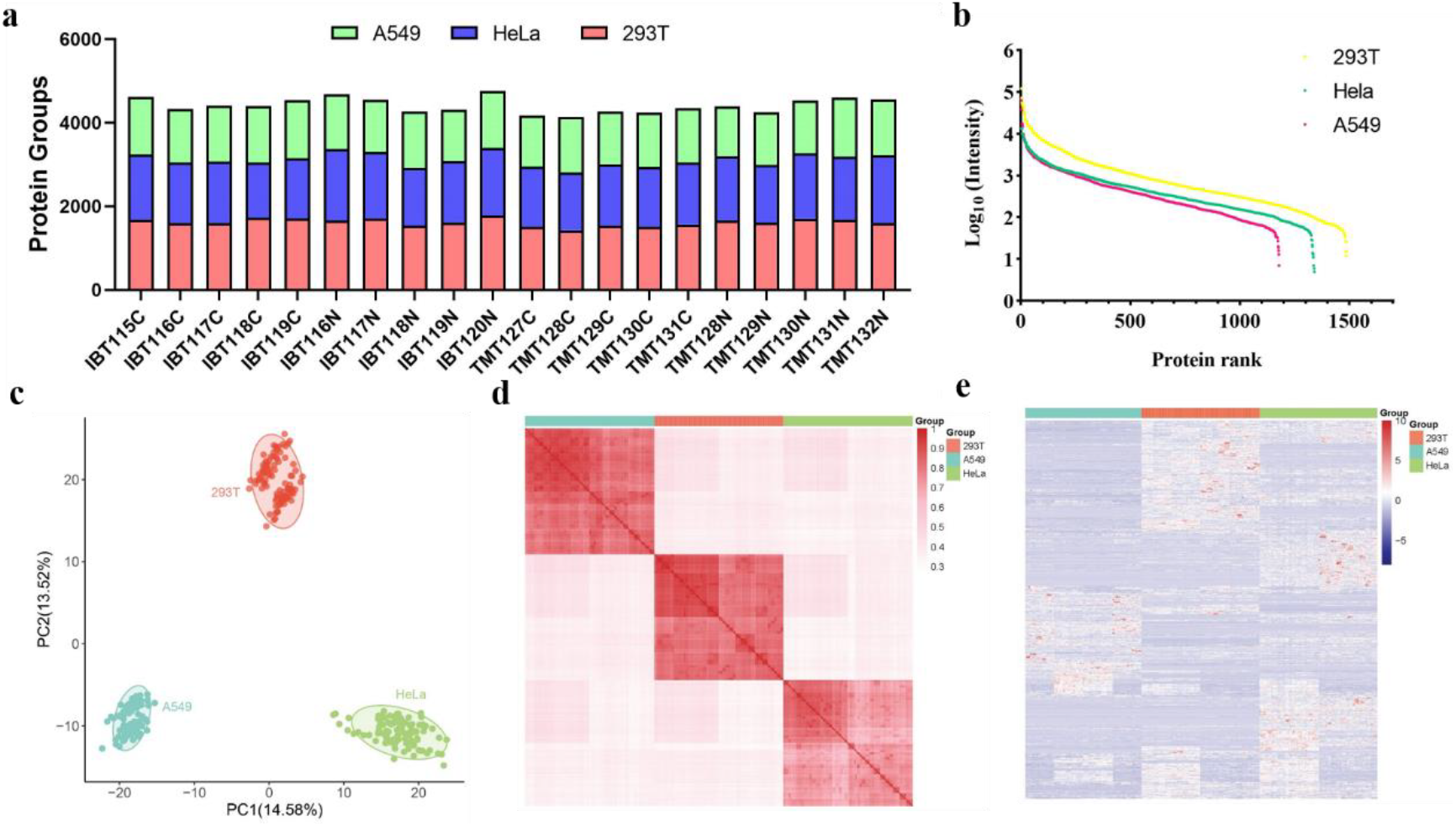
Evaluation of quantitative results from TMT-IBT-labelled 293T, HeLa, and A549 single cells BOXmini^TM^ SCP. (a) Number of quantified proteins, (b) Dynamic ranges (c) PCA analysis, (d) Pearson correlation analysis among three cell types, (e) Unsupervised hierarchical clustering of proteins.

Overall, we successfully balanced quantitative accuracy and proteome coverage in our ‘Integral-Hyperplex’ approach. Nearly 2,000 protein groups were accurately quantified from single cells, a number exceeding that of most published high-throughput single-cell proteomics methods. In this demonstration, throughput reached approximately 360 cells per day, primarily limited by the resolution of the timsTOF SCP, as only half of the available labelling channels were used. However, throughput can be further enhanced by adopting high-resolution, high-speed MS instruments such as Orbitrap Astral or timsTOF Ultra 2, and implementing shorter chromatography gradients. In theory, up to ∼2,000 single cells per day could be analyzed using these improvements.

## CONCLUSION

We propose a novel ‘Integral-Hyperplex’ strategy for high-throughput scProteomics by fully integrating IBT16 and TMT16 labelling within a single LC-MS/MS experiment. Because the signal responses of TMT16 and IBT16 reporter ions differ substantially, we developed two normalization methods for quantitative analysis: (1) employing a normalization channel in both IBT and TMT groups to serve as a ‘normalization standard’, and (2) applying protein-specific conversion factors to normalize IBT16 and TMT16 signals for each protein. The normalization channel approach offers higher quantification accuracy, whereas the conversion factor method provides slightly greater throughput. Furthermore, we automated the labelling and coupling process using the BOXmini™ SCP instrument and minimized labelling costs to ∼ 0.47 US Dollar/cell. Compared to manual labelling, the BOXmini™ SCP system enabled identification of up to 2,000 proteins per single cell and significantly reduced variation. All reactions were conducted within nanodroplets on AM16K chips integrated into the BOXmini™ SCP, resulting in improved labelling efficiency and reduced reagent consumption. The throughput of our TMT-IBT method can be further increased by incorporating additional types of labelling reagents.

## Author Contributions

Hui Zhang: Methodology, Investigation, Formal analysis, Writing - Original Draft. Qing Zeng: Methodology, Investigation. Kai Jin: Investigation. Yi Liu: Formal analysis. Chuanxi Huang: Formal analysis. Xing Liu: Investigation. Jing Yang: Supervision. Hanbin Ma: Supervision. Fuchu He: Supervision. Yun Yang: Conceptualization, Supervision, Writing - Original Draft, Writing – Review & Editing.

## ACKNOWLEDGMENTS

We thank Dr. Pei Jiang for assistance in data analysis. This work was supported by the Ministry of Science and Technology of the People’s Republic of China (Grant No. 2020YFE0202200), the Pre-study Project of Phronesis Medicine Large-scale Scientific Facility funded by Guangzhou Development District, and Guangzhou National Laboratory (Grant No. GZNL2024A03001), the National Natural Science Foundation of China (No. 62374102), the China Postdoctoral Science Foundation (2022M722338), the Natural Science Foundation of Shandong Province (ZR2024QC161), the Suzhou Basic Research Pilot Project (SSD2024023), and the Science and Technology Innovation Project of Foshan, Guangdong Province, China (No.1920001000047).

## Notes

The authors declare no competing financial interest.

