## Supplementary Material for "High-throughput Single-cell Proteomics Enabled by an Integrated Hyperplexing and Automatic Labelling Approach without Fractionation"

### Contents


**EXPERIMENTAL SECTION** **S3**

**Figure S1.** **S5**

**Figure S2.** **S6**

**Figure S3.** **S7**

**Figure S4.** **S8**

**Figure S5.** **S9**

**Figure S6.** **S10**

**Figure S7.** **S11**

#### EXPERIMENTAL SECTION

**Materials.** TMTpro 16-plex reagent sets, TMT 0 reagent sets, and 50% hydroxylamine were purchased from Thermo Fisher Scientific (USA). Isobaric tag 16-plex (IBT 16-plex) reagent sets were purchased from Nanjing Apollomics Biotech Inc. (China).

**Cell Culture.** HeLa, A549, and human embryonic kidney (HEK) 293T cells were cultured at 37 ℃ with 5% $CO_2$ using Dulbecco's Modified Eagle Medium (DMEM) with 10% (v/v) fetal bovine serum (FBS) and 1:1000 Penicillin/Streptomycin. The culture medium was refreshed daily.

**Optimization of TMT Labelling and IBT Labelling conditions.**

HeLa digest in either HEPES or TEAB buffer was mixed with 1.0, 1.5, 2.0, or 2.5 μL of 5 μg/μL TMT0, or 0.5, 1.0, or 1.5 μL of 5 μg/μL IBT2 (115N, 118C), and incubated at room temperature for 2 h. Excess labelling reagents were quenched by adding 5% hydroxylamine. For each analysis, 1 ng of labelled HeLa digest was injected.

For single-cell analysis, 1 μL of lysis and digestion buffer (0.2% n-dodecyl-β-D-maltoside (DDM), 100 mM HEPES, and 20 ng/μL trypsin) was dispensed into each well of a 384-well plate using the cellenONE® system. Individual HeLa cells were sorted into the wells and incubated at 37 °C with 85% humidity for 2 h. TMT0 or IBT2 (115N, 118C) reagents were added manually to each well, followed by incubation at 1,000 rpm and room temperature for 2 h. The reactions were quenched by adding 5% hydroxylamine. Three manually labelled 293T cells (TMT0 and IBT2) were combined and analyzed by nanoLC-MS/MS.

**Bulk Sample Preparation for Carrier and Normalization channels.**

Cells were detached using 0.05% Trypsin-EDTA treatment and underwent thorough washing with phosphate buffered saline (PBS) three times. Subsequently, lysis buffer was added, and lysis was facilitated by sonication. The composition of the lysis buffer included 1% sodium deoxycholate (SDC), 100 mM tris (2-carboxyethyl) phosphine hydrochloride (TCEP, pH=7), 40 mM 2-chloroacetamide (CAA), and 100 mM tris（hydroxymethyl）aminomethane hydrochloride (Tris HCl, pH=8.0). Protein concentration was measured using a Nanodrop One. Trypsin was added at a 1:30

enzyme-to-protein ratio, and digestion proceeded overnight at 37 °C. After digestion, peptides were acidified to pH <3 and desalted using a Waters Sep-Pak tC18 cartridge. For the preparation of carrier peptides, TMTpro-133C, TMTpro-134N, IBT16-121C, or IBT16-122 was used to label peptides. For normalization channels, peptides were labelled with one of the following labels: TMTpro-126, TMTpro-127N, IBT16-114, or IBT16-115N. Labelling reactions were quenched using 5% hydroxylamine.

**NanoLC-MS/MS analysis on timsTOF SCP.**

NanoLC-MS/MS analysis was conducted on a hybrid trapped ion mobility spectrometry (TIMS) quadrupole time-of-flight mass spectrometer (timsTOF SCP, Bruker Daltonics) along with a CaptiveSpray nano-electrospray ion source. Peptide desalting was carried out online using a 300 μm I.D. x 5 mm C18 PepMap100 trap column (Thermo Fisher Scientific, P/N 160454). Peptide separation was achieved on a nanoElute UHPLC system (Bruker Daltonics) coupled online to the timsTOF SCP via an integrated spray-tip analytical column (50 μm I.D. × 20 cm) packed with 1.9 μm/120 Å ReproSil-Pur C18 resin (Dr. Maisch GmbH). Mobile phases consisted of solvent A (water with 0.1% formic acid) and solvent B (acetonitrile with 0.1% formic acid). The flow rate was maintained at 100 nL/min, except for the first 5 min, during which it was increased to 200 nL/min. Peptide separation was performed using a 60-min gradient as follows: 5–8% B (0–5 min), 8–10% B (5–8 min), 10–26% B (8–47.6 min), 26–45% B (47.6–53 min), 45–95% B (53–55 min), and held at 95% B (55–60 min). Column temperature was maintained at 60 °C.

MS analysis was conducted in data-dependent acquisition (DDA) mode, scanning precursors over an m/z range of 100–1700. Ion mobility resolution ranged from 1.3 to 0.7 Vs $cm^{-2}$. Each acquisition cycle consisted of one full TIMS-MS scan followed by 10 parallel accumulation-serial fragmentation (PASEF) MS/MS scans. The ramp time was set to 166 ms, and precursors with charges from 0 to +5 were selected. Collision energy was stepped according to ion mobility, ranging from 44.80 eV at $1/K_0$ = 0.60 Vs $cm^{-2}$ to 89.60 eV at $1/K_0$ = 1.60 Vs $cm^{-2}$.

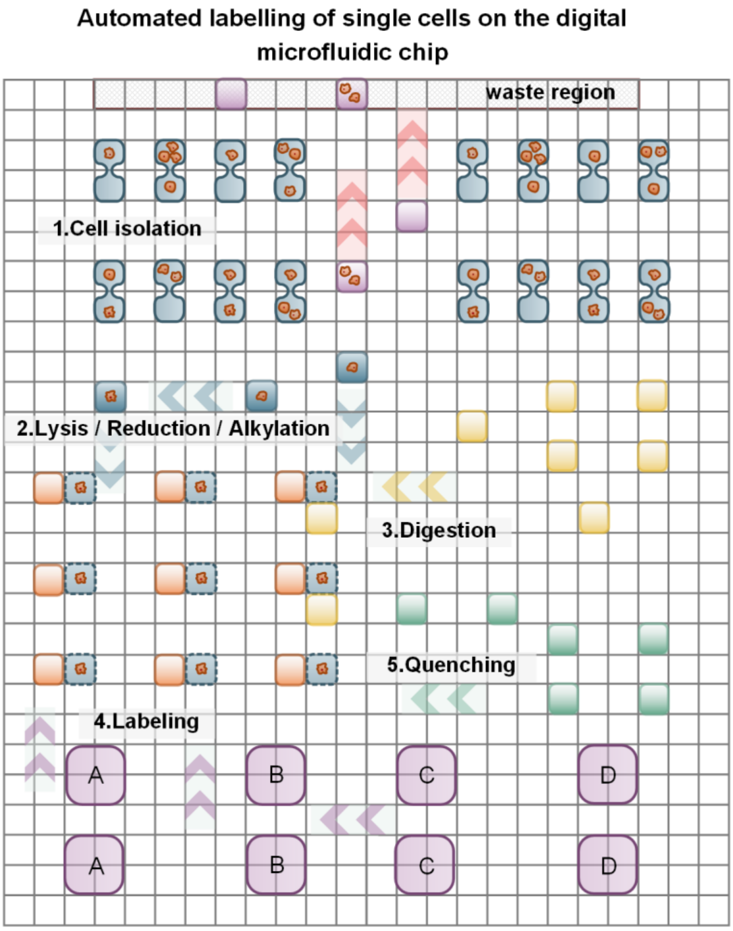


**Figure S1.** The expanded view of the layout and major reactions for automated labelling of single cells on the AM16K digital microfluidic chip.

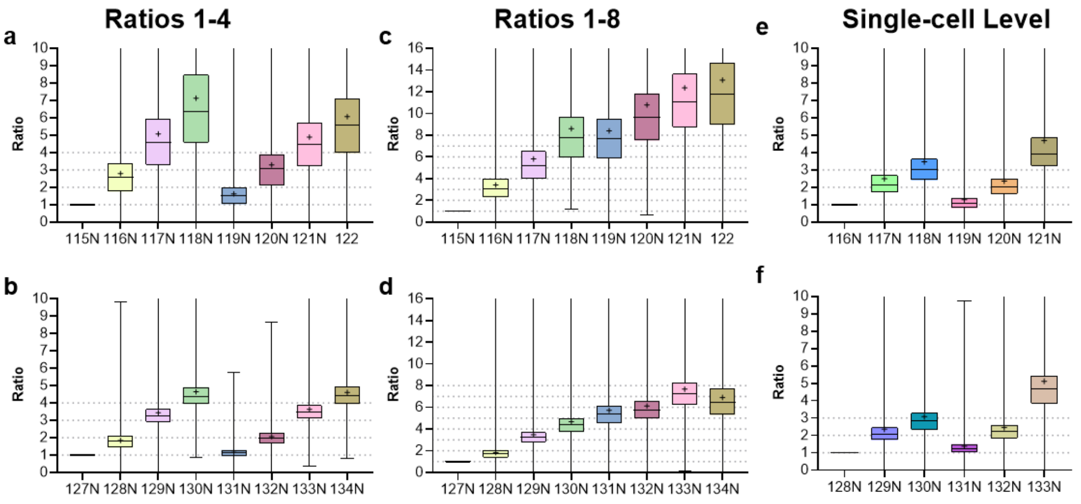


**Figure S2.** Evaluation of the quantitative accuracy of TMT and IBT labelling in Group B. (a-b) Proteins with a preset ratio of 1:2:3:4:1:2:3:4, labelled with (a) IBT16 or (b) TMT16. (c–d) Proteins with a preset ratio of 1:2:3:4:5:6:7:8, labelled with (c) IBT16 or (d) TMT16. (e–f) Observed ratios in IBT16 (e) and TMT16 (f) Group B, with a preset ratio of 1:2:3:1:2:3 (1 = 200 pg) and 50× carrier. Protein group ratios are plotted as box plots, with expected ratios represented by dashed lines. The "+" symbol represents the mean value and the "—" symbol indicates the median value.

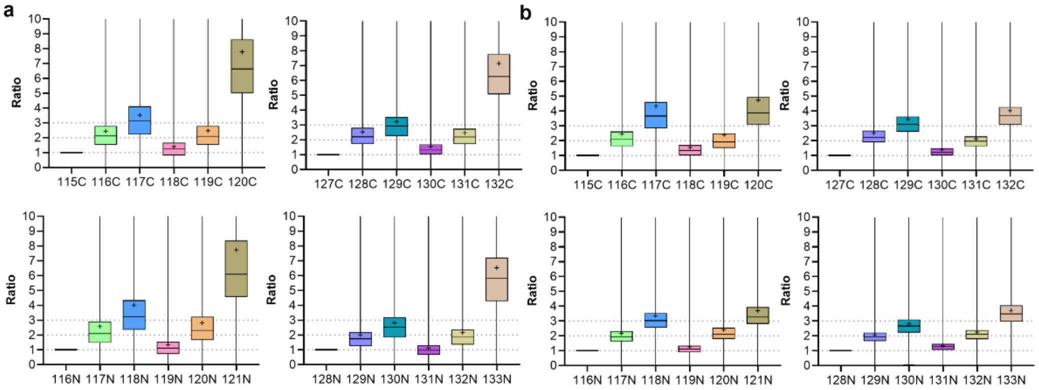


**Figure S3.** Evaluation of the quantitative accuracy with varying carrier dosages. Evaluation of quantitative accuracy at the single-cell-equivalent level with (a) 100× carrier or (b) 25× carrier. Experimental ratios were set to 1:2:3:1:2:3 for both.

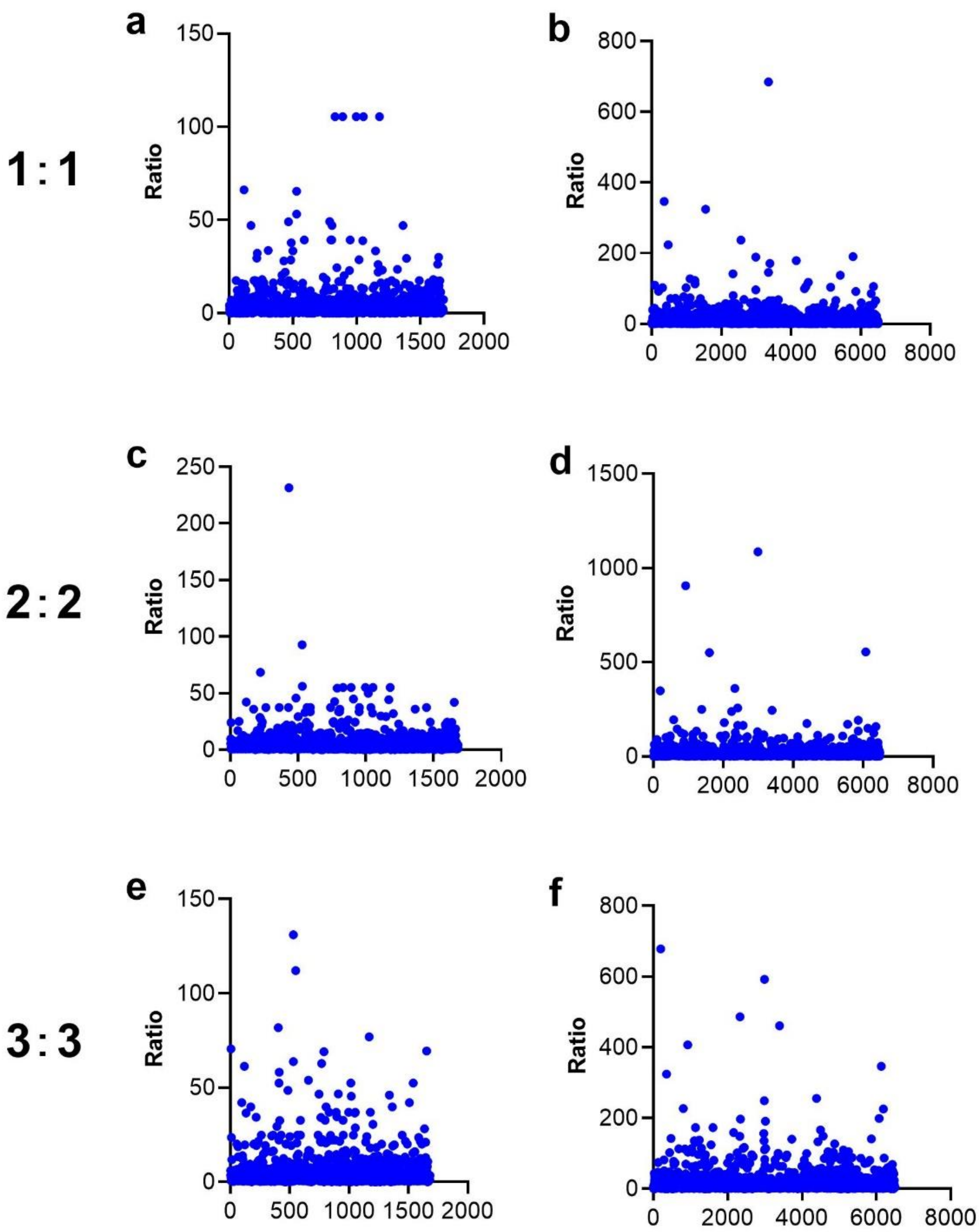


**Figure S4.** Quantitative differences for proteins and peptides between IBT and TMT groups. Scatter plots illustrate IBT/TMT ratios for proteins in the 1x channel (a), 2x channel (c) and (d) 3x channel, and for peptides in the 1x channel (a), 2x channel (c) and (d) 3x channel. Data are derived from Figure 4a.

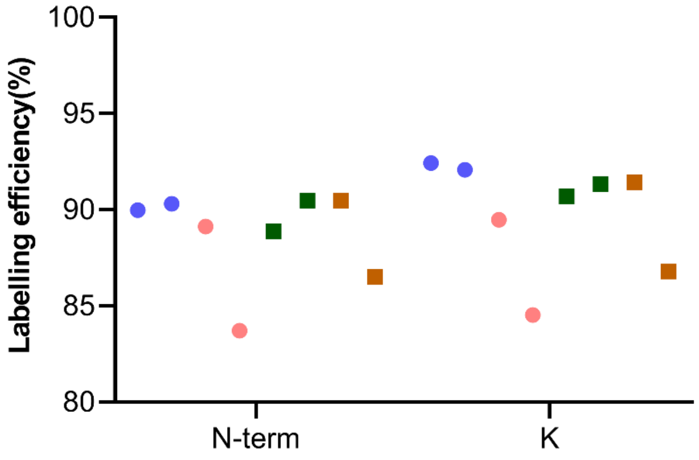


**Figure S5**. Labelling efficiency of manually labelled single 293T cells on peptide N-termini and lysine residues. Circles indicate IBT labelling, and squares indicate TMT labelling.

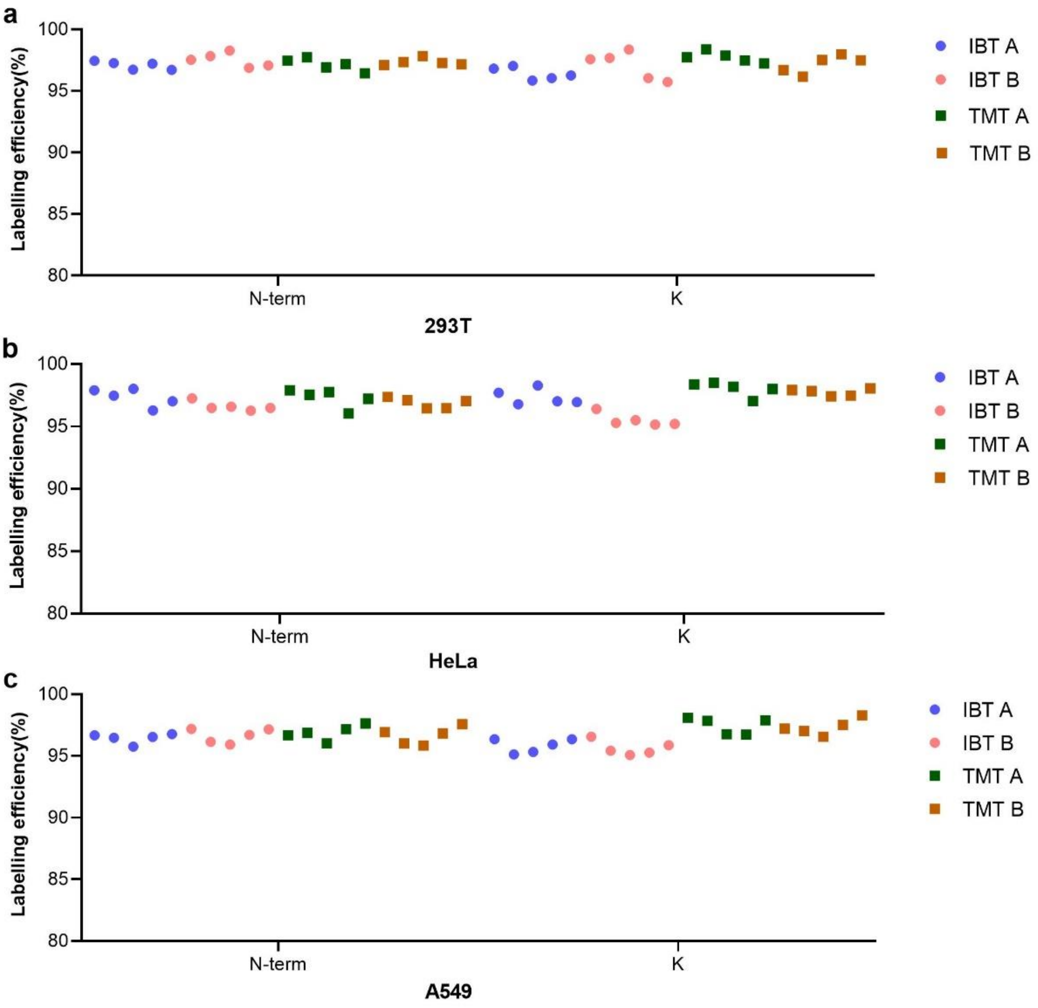


**Figure S6.** Labelling efficiencies of single cells labelled automatically using BOXmini™ SCP. Labelling efficiencies on peptide N-termini and lysine residues for (a) 293T, (b) HeLa, and (c) A549 cells. Circles indicate IBT labelling, and squares indicate TMT labelling.

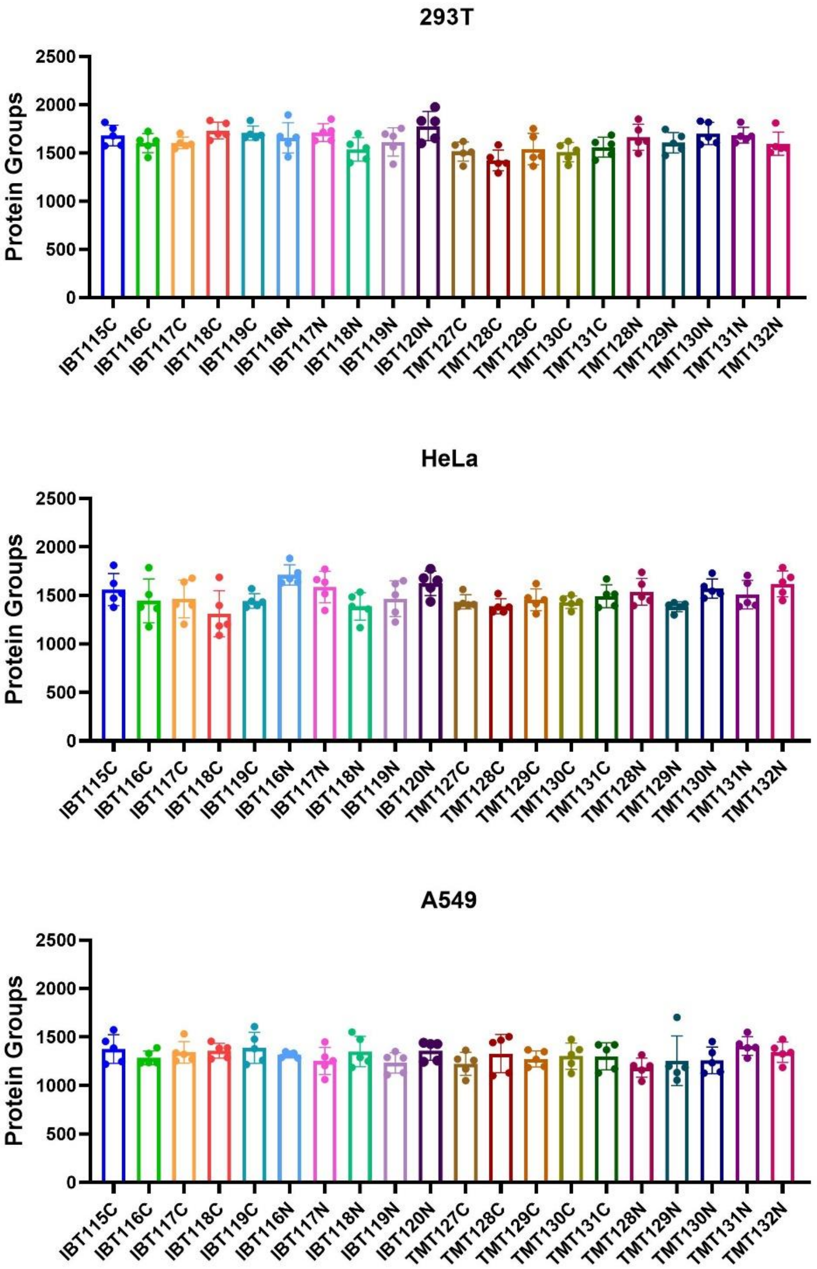


**Figure S7.** Labelling efficiencies of single cells labelled automatically using BOXmini™ SCP. Each cell type has 5 replicates.
